# Transcriptomic and physiological analyses reveal a salinity-induced growth suppression and defense activation in spring barley

**DOI:** 10.64898/2026.07.30.741898

**Authors:** Ammar Elakhdar, Elhamy Abdelwahab, Dina Elmoghazy, Takahiko Kubo

## Abstract

Salinity is a major abiotic stress that severely limits plant growth and productivity, causing substantial yield losses. Despite barley’s relative tolerance to salinity, the underlying physiological and molecular mechanisms remain incompletely understood. In this study, we employed an integrative approach combining agronomic, physiological, biochemical, and transcriptomic analyses to investigate salinity responses in the spring barley cultivar *Giza 134* under both field and lysimeter-based conditions. Salinity stress significantly reduced growth and yield-related traits, with more pronounced effects observed under lysimeter-imposed salinity, reflecting higher stress intensity. These reductions were associated with impaired water status, altered leaf structural traits, and declines in photosynthetic pigment content. In contrast, proline accumulation increased, indicating activation of osmotic adjustment mechanisms. Salinity also disrupted ionic homeostasis, as evidenced by elevated Na^+^ levels, reduced K^+^ content, and an increased Na^+^/K^+^ ratio. Enhanced lipid peroxidation and elevated catalase and peroxidase activities suggested increased oxidative stress and activation of antioxidant defenses. Transcriptome profiling identified 4,298 differentially expressed genes, including 1,764 upregulated and 2,534 downregulated genes. Functional enrichment analyses revealed upregulation of pathways related to stress adaptation, redox regulation, and metabolic reprogramming, while genes associated with photosynthesis, ribosome biogenesis, and protein synthesis were strongly suppressed. Several novel stress-responsive genes involved in signaling, osmoprotection, antioxidant defense, and central metabolism were highly induced, supported by coordinated enrichment of cis-regulatory motifs in their promoter regions.

Together, these findings provide a comprehensive physiological and molecular framework for salinity tolerance in Giza 134 and highlight candidate genes and pathways for breeding salt-resilient cultivars suited to saline-prone environments.

## 1. INTRODUCTION

Soil salinity is an increasingly serious global challenge and is widely recognized as one of the most detrimental abiotic stresses limiting crop productivity and threatening the long-term sustainability of agricultural systems. Currently, approximately 10.7% of the global land area, equivalent to around 1,381 million hectares, and nearly 20% of cultivated arable land are affected by varying degrees of salinization. This problem is projected to intensify further due to rising sea levels, unsustainable irrigation practices, and the accelerating impacts of climate change (FAO 2024; Rengasamy 2006). As global food demand continues to rise while arable land becomes increasingly salinized, the development of salt-tolerant crop cultivars has emerged as a critical priority, necessitating a deeper understanding of plant physiological and molecular responses to salinity stress.

Excessive soil salinity impairs plant water uptake and disrupts ionic homeostasis, primarily through the accumulation of toxic concentrations of sodium (Na^+^) and chloride (Cl^-^) ions. These conditions induce osmotic stress, nutrient imbalances, oxidative damage, and metabolic dysfunction, ultimately leading to reduced growth and yield (Tester and Davenport 2003; White 2001). To mitigate these effects, plants activate a complex array of physiological, biochemical, and molecular mechanisms, including osmotic adjustment, ion exclusion or compartmentalization, enhancement of antioxidant defenses, and modulation of growth and developmental processes (Munns and Tester 2008; Tester and Davenport 2003).

At the cellular and molecular levels, salinity stress triggers extensive reprogramming of gene expression, protein synthesis, and metabolic activity. Key genes involved in ion transport and homeostasis, such as *SOS1, HKT1,* and *NHX,* play central roles in maintaining cellular ion balance (Apse et al. 1999; Rivandi et al. 2011). In parallel, pathways associated with osmolyte biosynthesis (e.g., proline and glycine betaine), reactive oxygen species (ROS) scavenging enzymes (e.g., SOD, CAT, and POD), and hormonal signaling networks, including abscisic acid (ABA), ethylene, and jasmonic acid contribute to stress mitigation and adaptive responses (Ashraf and Foolad 2007; Elsawy et al. 2018; Wyn Jones and Storey 1978) (Riemann et al. 2015; Sreenivasulu et al. 2006). Transcription factors (TFs) act as key regulators of these stress-responsive gene networks, with families such as AP2/ERF, bZIP, DREB, MYB, NAC, WRKY, and HSF orchestrating the activation or repression of downstream targets during salinity stress (Alexander et al. 2019; Elakhdar et al. 2023; Guerin et al. 2019; Guo et al. 2016). Phytohormones, particularly ABA, further fine-tune adaptive responses by regulating stomatal conductance, osmotic balance, and stress-induced gene expression (Socias et al. 1997). Despite significant progress, the complex, multigenic, and developmentally dynamic nature of salinity tolerance remains incompletely understood. Barley (*Hordeum vulgare L*.), one of the earliest domesticated cereal crops, serves as a valuable model for studying abiotic stress tolerance due to its diploid genome, well-annotated reference sequences, and considerable natural variation in stress responses (Coulter et al. 2022; International Barley Genome Sequencing et al. 2012; Mascher et al. 2017; Schulte et al. 2009). Compared with wheat and rice, barley generally exhibits greater salinity tolerance and is widely cultivated in marginal and salt-affected environments. Nevertheless, salinity stress continues to cause substantial yield losses, particularly during early developmental stages, underscoring the need for further genetic improvement (Colmer et al. 2005; Munns et al. 2006).

Recent advances in high-throughput sequencing technologies have accelerated the dissection of complex traits such as salinity tolerance. RNA sequencing (RNA-seq) enables genome-wide characterization of transcriptional responses to stress, providing insights into regulatory pathways and biological processes underlying plant adaptation (Elakhdar et al. 2023; Mutz et al. 2013). Comparative transcriptomic approaches facilitate the identification of differentially expressed genes (DEGs) across genotypes, developmental stages, or environmental conditions, thereby enabling the discovery of key regulatory modules and candidate genes involved in stress perception, signal transduction, and protective responses.

Although several studies have investigated barley responses to short-term salinity stress, particularly at the seedling stage, comprehensive analyses of prolonged salinity exposure across multiple developmental stages remain limited. Integrating transcriptomic data with detailed physiological and biochemical measurements is therefore essential for achieving a holistic understanding of salinity tolerance mechanisms. In this study, we employed an integrative approach to investigate the salinity stress response of the spring barley cultivar Giza 134. Morphological, physiological and biochemical measurements were evaluated under salt stress at both the seedling and adult stages, in addition to RNA-seq analysis was conducted to capture genome-wide transcriptional changes. This comprehensive analysis enabled the identification of key salinity-responsive genes in Giza 134 and regulatory pathways, offering valuable insights for breeding and biotechnological strategies aimed at enhancing salinity tolerance in barley.

## 2. Materials and methods

### 2.1. Plant materials

Two independent experiments were performed to evaluate the response of the spring barley (*Hordeum vulgare* L.) cultivar ‘Giza 134’ (Alanda-01/4/WI2291/3/Api/CM67//L2966-69) to salinity stress at seedling and adult growth stages. Giza 134 is a high-yielding cultivar recommended for cultivation in newly reclaimed lands in Egypt, with an average grain yield of approximately 4600 kg ha^-1^. It exhibits broad resistance to major foliar diseases, including leaf rust, stripe rust, and powdery mildew. Due to its consistent performance under abiotic stress conditions, Giza 134 serves recently as a robust genetic model for dissecting salinity-and drought-responsive mechanisms at the transcriptomic level (Elakhdar et al. 2023). Certified seeds were kindly provided by the Barley Research Department, Field Crops Research Institute, Agricultural Research Center (ARC), Egypt.

### 2.2. Field salinity experiment

Field trials (SS1) were conducted over two consecutive growing seasons (2023/2024 and 2024/2025) at two experimental sites of the ARC, to evaluate the effect of natural soil salinity on yield components of the spring barley cultivar Giza 134. The two locations were Sakha Agricultural Research Station and El-Husseiniya Agricultural Research Station, representing normal and saline-affected soils, respectively.

### 2.3. Lysimeter-based field system

A lysimeter-based system was established to monitor soil salinity dynamics and plants responces under salinity condtion. The system comprised 48 rectangular lysimeters (2.3 m × 1.8 m × 1.2 m; length × width × depth) embedded within a drainage pond. Each lysimeter was internally lined with fired ceramic tiles to prevent lateral water movement. Saline irrigation water was stored in overhead tanks and delivered via a submersible pump through surface emitters. Drainage water was collected using perforated PVC pipes wrapped in jute fabric to prevent clogging and facilitate leachate sampling. Plants were subjected to two irrigation regimes: Well-watered (N): irrigation with fresh water, and salinity stress (SS2): irrigation with saline water (16 dS m^-1^ NaCl). The experiment followed a randomized complete block design with three replicates. Drip irrigation ensured uniform water distribution. At physiological maturity, samples were harvested from the central area of each plot (2 m^2^), and biological and grain yields were recorded.

### 2.4. Growth chamber experiment

To investigate transcriptomic responses to salinity stress, a growth chamber experiment was conducted at the Institute of Plant Genetic Resources, Kyushu University (Ito Campus, Fukuoka, Japan). Seeds of Giza 134 were surface-sterilized in 0.5% (v/v) sodium hypochlorite (NaOCl) containing Tween 20 (Sigma-Aldrich) for 15 minutes, followed by thorough rinsing with distilled water. Sterilized seeds were sown in plastic pots (23 × 23 × 19 cm) filled with commercial seedling-raising soil (JA Kumiai King Soil; Agr. Japan Co., Ltd.), with four plants per pot. Pots were kept in the dark at 25 °C for three days to promote coleoptile emergence. Subsequently, seedlings were transferred to a growth chamber (SANYO) maintained at 27 °C during a 14-hour light period and 25 °C during a 10-hour dark period. Soil moisture was maintained at 40–50% of field water capacity (FWC) for 10 days prior to stress application. The experiment followed a completely randomized design with three biological replicates, each consisting of 12 seedlings. Plants were assigned to either a well-watered, irrigated with tap water, or a salinity stress treatment, irrigated with NaCl solution at an electrical conductivity of 16 dS m^-1^ for 15 days. At the end of the treatment period, shoot tissues from three plants per treatment were harvested, immediately frozen in liquid nitrogen, and stored at −80 °C for RNA extraction.

### 2.5. Trait measurements

Phenotypic, agronomic, and physiological traits were evaluated under both irrigation regimes.

#### 2.5.1. Relative water content (RWC) determination

Leaf relative water content (RWC, %) was determined using:

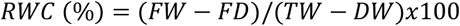

where FW = fresh weight, TW = turgid weight after 4 h hydration at 25 °C, and DW = dry weight after oven drying at 70 °C.

#### 2.5.2. Leaf Morphological Traits

Fully expanded leaves were used to measure:

Leaf morphological traits, including leaf area (LA), specific leaf area (SLA), and specific leaf weight (SLW), were determined using fully expanded leaves.

##### Leaf Area (LA)

Leaf area (LA; cm^2^) was measured using a leaf area meter and expressed as cm^2^

##### Specific Leaf Area (SLA)

Specific leaf area (SLA; cm^2^ g^-1^) was calculated as the ratio of leaf area to leaf dry weight using the following equation:

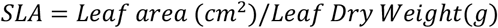

##### Specific Leaf Weight (SLW)

Specific leaf weight (SLW; g cm^-2^), representing leaf thickness and density, was calculated as the inverse of SLA:

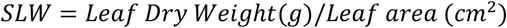

Leaves were oven-dried at 70 °C until constant weight before determining dry mass.

#### 2.5.3. Na^+^ and K^+^ Determination

Dried leaf tissues were ground (LM-Plus grinder, Osaka Chemical Co.) and digested using H_2_SO_4_:H_2_O_2_ (2:1, v/v). Na^+^ and K^+^ ion concentrations were measured using a flame photometer (ANA-135, Tokyo Photoelectric, Japan).

#### 2.5.4. Antioxidant Enzyme Assays

Antioxidant enzyme activities were determined to evaluate the oxidative stress response of barley plants under salinity conditions. Fresh leaf tissue (0.5 g) was immediately frozen in liquid nitrogen and homogenized using a chilled mortar and pestle in 10 mL of ice-cold phosphate buffer (0.2 M, pH 7.0). The homogenate was centrifuged at 15,000 × g for 10 min at 4 °C, and the resulting supernatant was collected and used as a crude enzyme extract for subsequent enzymatic assays.

Catalase (CAT; EC 1.11.1.6) was measured following the method of (Kato and Shimizu 1987), based on the decomposition of hydrogen peroxide (H_2_O_2_). The reaction mixture, with a final volume of 3 mL, consisted of 0.5 mL phosphate buffer (0.2 M, pH 7.0), 0.3 mL H_2_O_2_ solution, and 0.1 mL enzyme extract, with the remaining volume adjusted using distilled water. The decrease in absorbance resulting from H_2_O_2_ degradation was monitored at 240 nm using a UV–Vis spectrophotometer. Catalase activity was expressed as µmol H_2_O_2_ decomposed min^-1^ g^-1^ protein.

Peroxidase (POD; EC 1.11.1.7) was determined according to the method of (Chance and Maehly 1955), which measures the oxidation of guaiacol in the presence of hydrogen peroxide. The reaction mixture contained 2.5 mL phosphate buffer (0.1 M, pH 6.0), 0.3 mL guaiacol solution, 0.1 mL H_2_O_2_, and 0.1 mL enzyme extract. The increase in absorbance caused by the formation of tetraguaiacol was recorded at 470 nm for 1 min using a spectrophotometer. Peroxidase activity was expressed as the change in absorbance per minute per gram fresh weight.

#### 2.5.5. Lipid peroxidation (MDA Content)

Lipid peroxidation was assessed by measuring malondialdehyde (MDA) content following the method of (Heath and Packer 1968). Fresh leaf tissue (0.1 g) was homogenized in 0.5 mL of 0.1% (w/v) trichloroacetic acid (TCA) and centrifuged at 15,000 × g for 10 min at 4 °C. The supernatant (0.5 mL) was mixed with 1.5 mL of 0.5% thiobarbituric acid in 20% TCA and incubated at 95 °C for 25 min. After cooling, absorbance was measured at 532 and 600 nm. MDA content was expressed as µmol g^-1^ fresh weight.

#### 2.5.6. Chlorophyll Determination

Chlorophyll a and b were determined following Moran (Moran 1982). Briefly, fresh leaf tissues were homogenized in *N-N-*dimethylformamide (C_3_H_7_NO) and incubated overnight at 4 °C to ensure complete pigment extraction. Absorbance was measured at 664 nm and 647 nm using a spectrophotometer. Chlorophyll concentrations were calculated using the equations described by Lichtenthaler (Lichtenthaler 1987).

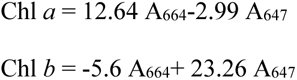

where A_664_ and A_647_ are the absorbance at wavelengths 664 and 647, respectively.

#### 2.5.7. Proline Content

Proline accumulation was quantified according to (Bates et al. 1973). Leaf tissue (0.5 g) was homogenized in 10 ml of 3% (w/v) sulfosalicylic acid (C_7_H_6_O_6_S) and filtered through Whatman No. 1 filter paper. Filtrate (2 mL) was reacted with 2 mL glacial acetic acid (CH_3_COOH) and 2 mL acid ninhydrin (C_9_H_6_O_4_) and incubated at 100 °C for 1 h. After cooling, 4 mL toluene (C_6_H_5_CH_3_) was added, and the mixture was gently shaken to separate the chromophore-containing toluene layer. The absorbance of the chromophore-containing phase was measured at 520 nm. Proline content was expressed as mg g^-1^ fresh weight.

#### 2.5.8. Agronomic Traits

Agronomic traits evaluated included plant height (PH; cm), Biology yield (BY; kg), Grain yield (GY: kg), and 1000 grain weight (1000 GW).

#### 2.5.9. Statistical Analysis

Differences between treatments were determined using Student’s t-test. All analyses were performed using three biological replicates, and statistical significance was determined at *P* < 0.05.

### 2.6. RNA isolation and high-throughput sequencing

Total RNA was extracted from 100 mg of leaf blade tissue using the RNeasy® Plant Mini Kit (QIAGEN) following the manufacturer’s protocol. RNA quantity and purity were assessed using a NanoDrop 2000 spectrophotometer (Thermo Scientific), and RNA integrity was verified before library construction. Paired-end cDNA libraries (150 bp) were prepared and sequenced using the DNBSEQ platform, following the manufacturer’s instructions.

### 2.7. Mapping and transcriptome analysis

Raw sequencing reads were first quality-checked and filtered to remove low-quality sequences and adapter contamination using Cutadapt (Martin 2011). Clean reads were aligned to the barley reference genome (IBSC_v2) using HISAT2 (*v2.2.0*) (Kim et al. 2015). Transcript assembly and quantification of aligned reads were performed using StringTie (*v2.2.1*) (Pertea et al. 2016). Differential gene expression (DEGs) analysis between normal and salinity stress treatments was conducted using DESeq2 (Love et al. 2014). Significantly DEGs were filtered based on FDR-adjusted *p-*value ≤0.05 with log2 FC ≥1.5 for further analysis. Gene annotation was performed using BARLEYMAP (Cantalapiedra et al. 2015), the Morex Genome Map (Mascher et al. 2017), and the James Hutton iSelect annotation database (https://ics.hutton.ac.uk/50k/). Functional annotation was supported by BLAST and position-specific scoring matrices (PSSMs).

### 2.8. Functional annotation, gene ontology, and pathway enrichment analyses

Gene Ontology (GO) enrichment analysis of the DEGs was conducted using the ShinyGO platform (Ge et al. 2020), with a significance threshold of *P* ≤ 0.05 and FDR ≤ 0.05. DEGs exhibiting consistent expression changes (either upregulated or downregulated) were grouped based on their expression patterns. To gain deeper insight into the metabolic pathways associated with the DEGs, Kyoto Encyclopedia of Genes and Genomes (KEGG) pathway enrichment analysis was performed using the PlantGSAD (Ma et al. 2022). Hierarchical clustering of gene expression data was performed using the *pheatmap* package in R (*v1.0.12*). Additionally, volcano plots were generated using GraphBio (Zhao and Wang 2022) to illustrate the overall distribution of gene expression changes.

### 2.9. Identification of *cis*-regulatory elements in promoter regions

To identify *cis*-regulatory motifs associated with transcriptional start sites (TSS), we analyzed the 1 kb upstream regions (−1000 to −1 bp upstream of the start codon) of 1,764 upregulated and 2534 downregulated genes. Overrepresented hexamer oligonucleotides and positional sequence patterns were identified using RSAT motif discovery tools based on a Markov chain model (Turatsinze et al. 2008). Promoter sequences from the barley reference genome (*Hordeum vulgare L.*) were retrieved with the “retrieve sequence” tool in RSAT. The positional enrichment of motifs was evaluated by calculating χ^2^-based *P-*values, comparing the observed positional distribution against the expected distribution under a homogeneous model (assuming equal probability of oligonucleotide occurrence across the region). For motif annotation, the TOMTOM comparison tool in the MEME Suite (https://meme-suite.org) was used to identify homologous matches to known transcription factor (TF) binding motifs against the JASPAR CORE plant and *Arabidopsis* databases.

### 2.10. Tissue-specific expression profile

Expression profiles of 40 candidate genes across 16 barley tissues and developmental stages were retrieved from the BarleyNBI functional omics platform (www.inetbio.org/barleynet/). The data values are presented as normalized Transcripts Per Million (TPM). Expression values were obtained as normalized Transcripts Per Million (TPM). The TPM matrix was downloaded and processed in R. TPM values were log_2_-transformed [log_2_(TPM + 1)], standardized using z-scores for each gene to highlight tissue-specific expression patterns, and visualized using *pheatmap* with Euclidean distance and complete linkage clustering. The resulting heatmap represents standardized expression values (z-scores), where blue indicates expression below the mean and red indicates expression above the mean.

### 2.11. cDNA library construction and gene validation

First-strand cDNA was synthesized from 1.5 µg total RNA. Reverse was performed using the PrimeScript™ RT reagent kit (TaKaRa), following the protocol described (Elakhdar et al. 2023). Expression levels of salinity-related genes were quantified in three technical replicates. RT-qPCR was performed using the KAPA SYBR® FAST qPCR kit (Kapa Biosystems) on a CFX96 Real-Time PCR System (Bio-Rad). Cycling conditions were 95 °C for 60 s followed by 40 cycles of 95 °C for 15 s and 60 °C for 60 s. *β-TUBULIN* was used as an internal reference gene, and relative expression levels were calculated using the comparative Ct method (2^ΔΔCt^) (Schmittgen and Livak 2008). Gene-specific oligonucleotide primers were designed using the Primer3 tool via NCBI Primer-BLAST. Primer sequences are provided in Table S1.

## 3. Results

### 3.1. Growth Performance and Agronomic Responses

Salinity stress significantly affected the growth performance and agronomic productivity of the barley cultivar *Giza 134* (Figure 1). Compared with the normal (N), both field salinity (SS1) and lysimeter-based salinity stress (SS2) resulted in significant reductions in plant height (PH), biological yield (BY), grain yield (GY), and 1000-grain weight (1000 GW) (Figure 1). Plant height decreased by approximately 15–20% under SS1 and by nearly 25% under SS2 relative to N conditions (*P* < 0.01). A comparable pattern was observed for biological yield, which declined significantly under both salinity treatments, with a more pronounced reduction under SS2 (*P* < 0.001). Grain yield exhibited the highest sensitivity to salinity stress, showing a significant reduction under SS1 and a further decline under SS2, indicating the stronger inhibitory effect of lysimeter-imposed salinity conditions. Similarly, 1000-grain weight was significantly reduced under both salinity regimes, with the largest decrease recorded under SS2 (*P* < 0.001).

**Figure 1.**
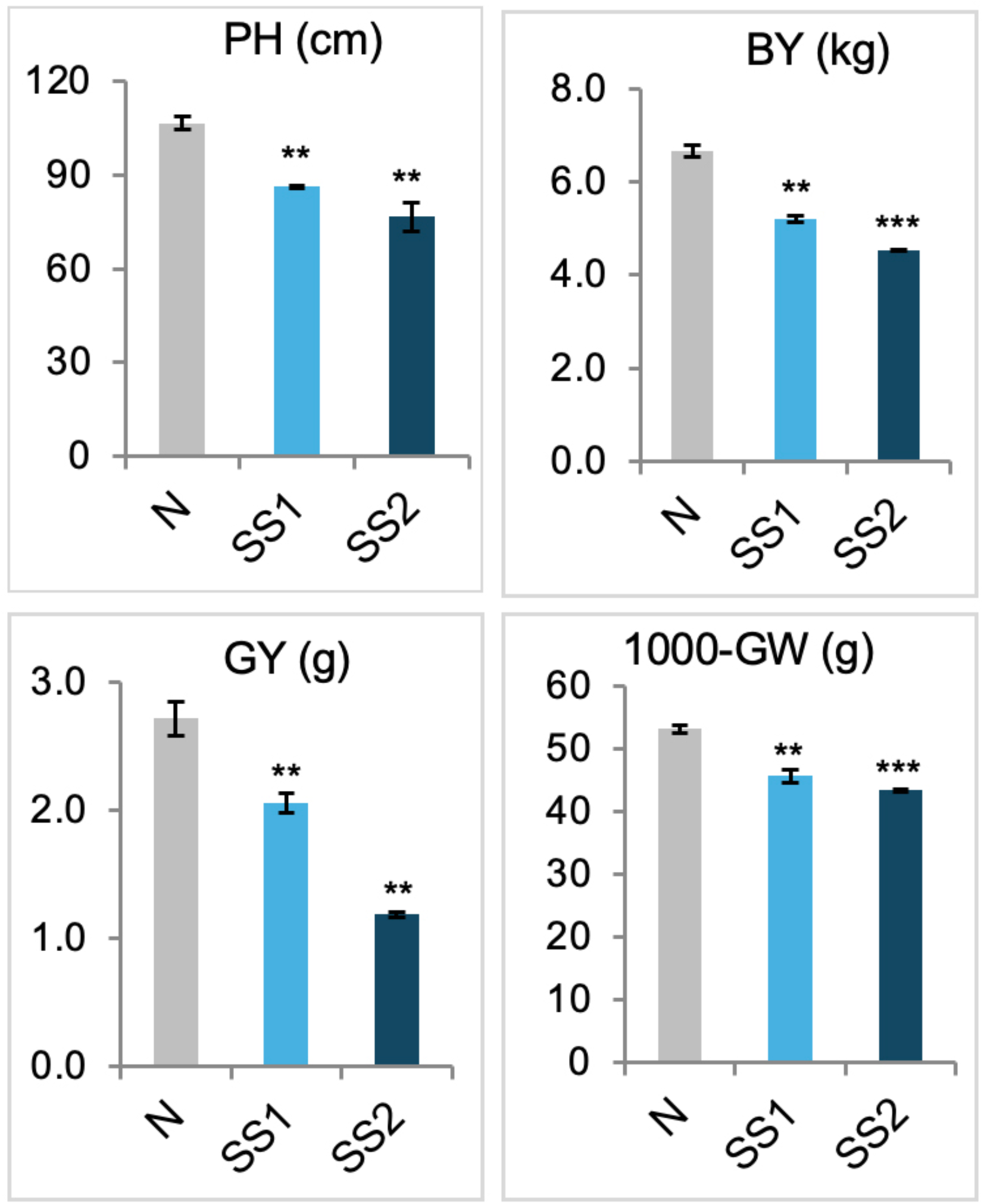
Agronomic performance of the spring barley cultivar Giza 134 under salinity stress. Plant height (PH, cm), biological yield (BY, kg), grain yield (GY, kg), and 1000-grain weight (1000-GW, g) measured under control and salinity conditions. Values are presented as mean ± standard error (SE) of three biological replicates. Asterisks indicate statistically significant differences between treatments according to Student’s *t-test* (*P* < 0.05).

### 3.2. Leaf Morphological and Plant Water Status Responses to Salinity Stress

Salinity stress significantly affected leaf morphological traits and plant water status in barley (Figure 2a). Leaf area (LA) decreased significantly under saline conditions compared with the well-watered control (*P* < 0.01), indicating reduced leaf expansion under stress. Consistent with this trend, specific leaf weight (SLW) decreased markedly under saline conditions (*P* < 0.001), reflecting reduced leaf thickness and biomass accumulation per unit leaf area. These results demonstrate that salinity stress induced clear structural adjustments in leaf morphology.

**Figure 2.**
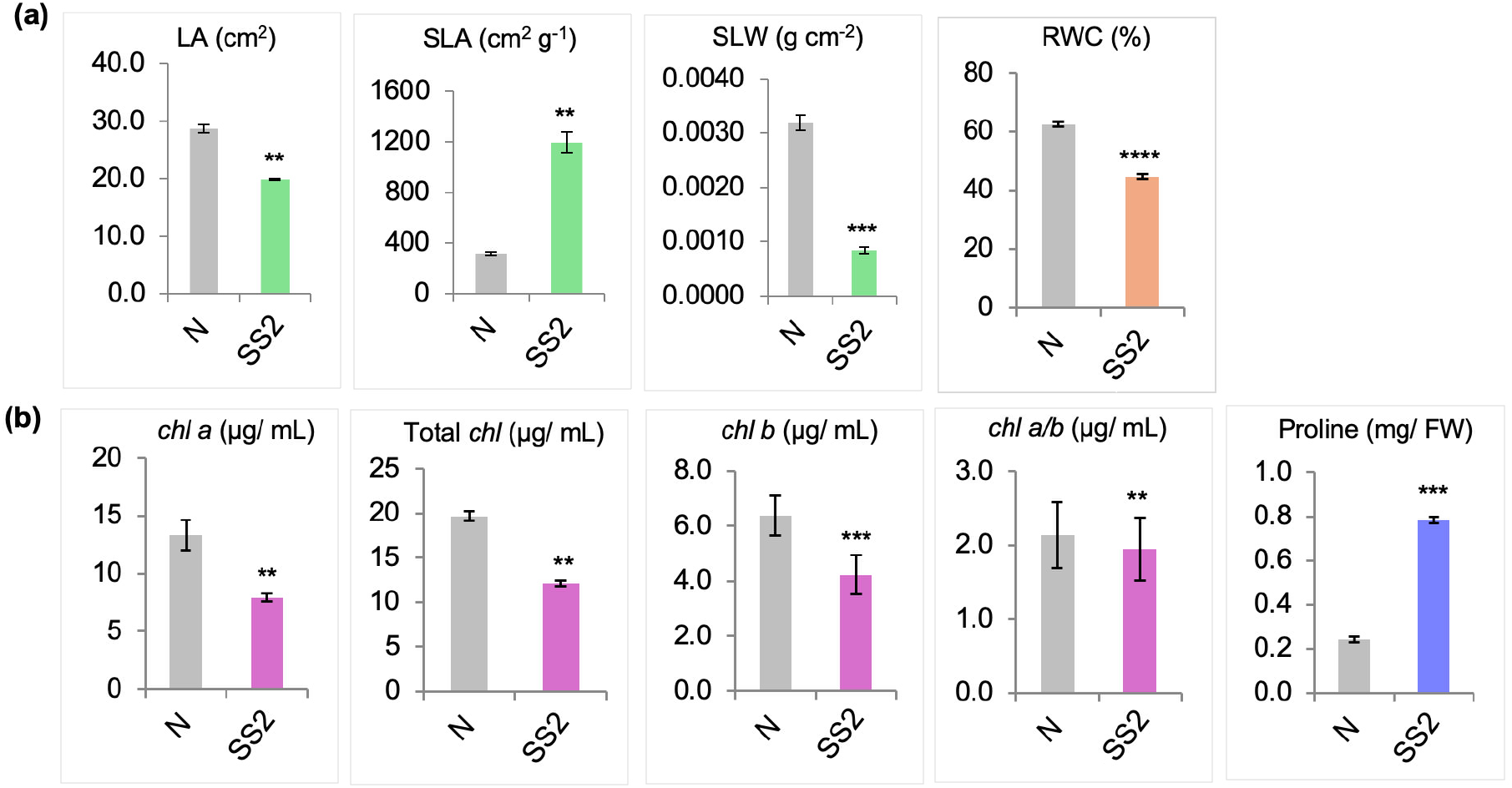
Morphological, physiological, and biochemical responses of the spring barley cultivar Giza 134 under salinity stress. (A) Leaf morphological traits, including leaf area (LA, cm^2^), specific leaf area (SLA, cm^2^ g^-1^), and specific leaf weight (SLW, g cm^-2^). Plant water status indicated by leaf relative water content (RWC, %). (B) Photosynthetic pigment and osmotic adjustment parameters, including chlorophyll *a*, chlorophyll *b*, chlorophyll *a/b*ratio, total chlorophyll content, and proline accumulation (mg g^-1^ FW). Values represent mean ± standard error (SE) of three biological replicates. Asterisks indicate statistically significant differences among different conditions by Student’s *t*-test.

Consistent with these morphological changes, relative water content (RWC) declined markedly under salinity stress (*P* < 0.0001), reflecting impaired plant water status and osmotic imbalance induced by saline conditions.

### 3.3. Photosynthetic Pigments and Osmotic Adjustment Responses

Salinity stress significantly influenced photosynthetic pigment composition and osmotic regulation (Figure 2b). Chlorophyll a content decreased significantly (*P* < 0.01), whereas chlorophyll b exhibited a stronger reduction (*P < 0.001*). Consequently, total chlorophyll content (Chl a + b) declined significantly (*P* < 0.01) under salinity stress. The chlorophyll a/b ratio also decreased significantly (*P* < 0.01), indicating alterations in pigment balance and potential impairment of photosynthetic efficiency.

In contrast, proline content increased markedly under salinity stress relative to the control treatment (*P* < 0.001). The substantial elevation of proline content suggests activation of osmotic adjustment mechanisms in response to salt-induced stress.

### 3.4. Ionic Homeostasis and Oxidative Stress Responses

Salinity stress caused substantial changes in ionic balance and oxidative stress markers. Sodium (Na^+^) concentration increased significantly (*P < 0.05*), whereas potassium (K^+^) content decreased significantly (*P < 0.05*). As a consequence, the Na^+^/K^+^ ratio increased markedly (*P* < 0.001) (Figure 3a), indicating pronounced ionic imbalance under saline conditions.

**Figure 3.**
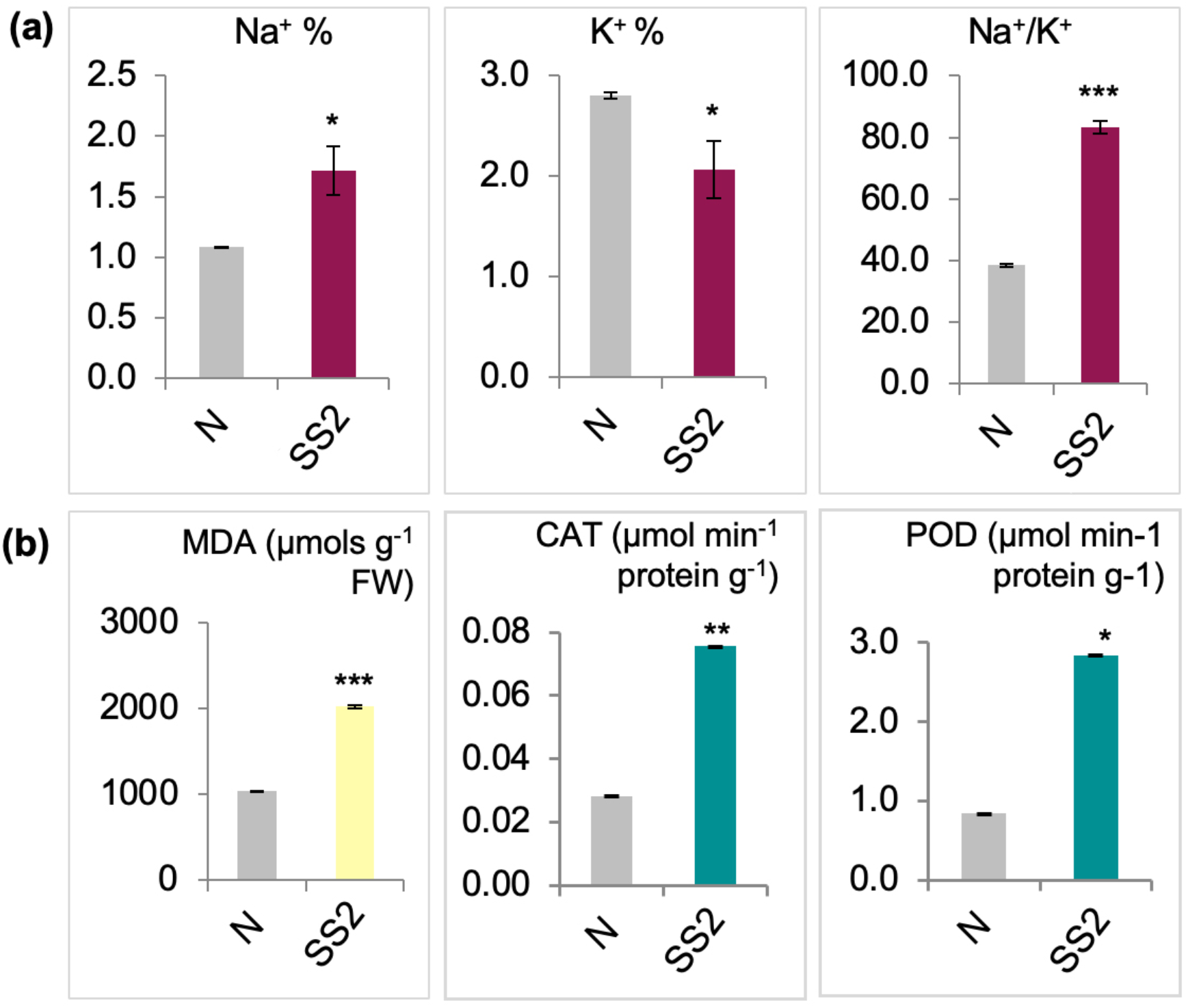
Ionic homeostasis and oxidative stress responses of the spring barley cultivar Giza 134 under salinity stress. (A) Sodium (Na^+^) content, potassium (K^+^) content, and Na^+^/K^+^ ratio under control and salinity conditions. (B) Antioxidant enzyme activities and lipid peroxidation markers, including catalase (CAT; µmol min^-1^ g^-1^ protein), peroxidase (POD; µmol min^-1^ g^-1^ protein), and malondialdehyde (MDA; µmol g^-1^ FW). Values are presented as mean ± standard error (SE) of three biological replicates. Statistical significance between treatments was determined using Student’s *t*-test (*P* < 0.05).

Lipid peroxidation, estimated by MDA content, increased significantly under salinity stress (*P* < 0.001) compared with control plants (Figure 3b), indicating enhanced membrane oxidative damage induced by salt stress.

Activities of antioxidant enzymes were significantly enhanced in response to salinity stress. CAT activity increased significantly (*P* < 0.01), while POD activity showed a moderate but significant increase (*P < 0.05*) compared with the well-watered treatment (Figure 3b). These results indicate activation of enzymatic antioxidant defense mechanisms under saline conditions.

### 3.5. Transcriptome analysis of salinity stress responses

To characterize transcriptional responses to salinity stress in Giza 134, RNA sequencing was performed on leaf tissues from WW and salinity-stressed (SS) plants. Sequencing was performed using the DNBSEQ platform, generating approximately 15.2 GB of raw data per lane. After quality filtering and adapter removal, high-quality clean reads were retained for downstream analyses

On average, 161.2 million (99.50%) and 127.7 million (99.62%) clean reads were obtained from WW and SS samples, respectively. More than 90% of reads mapped uniquely to the barley reference genome (*IBSC_v2*), indicating high sequencing quality and strong genome coverage suitable for transcriptome-wide analysis. Gene prediction and functional annotation using BARLEYMAP and the Morex Genome Map databases identified 23,991 expressed genes, providing a robust basis for differential expression analysis.

Differential expression analysis identified 4,298 differentially expressed genes (DEGs) in response to salinity stress, including 1,764 upregulated and 2,534 downregulated genes (Figure 4). The volcano plot revealed numerous genes exhibiting strong transcriptional responses, including several exhibiting more than 15-fold changes in expression (Figure 4a). The MA plot showed a broad and balanced distribution of expression changes across the transcriptome (Figure 4b), while the Venn diagram highlighted both unique and shared DEGs between control and salinity-stressed conditions (Figure 4c). Hierarchical clustering demonstrated clear separation between WW and SS samples and high reproducibility among biological replicates (Figure 4d).

**Figure 4.**
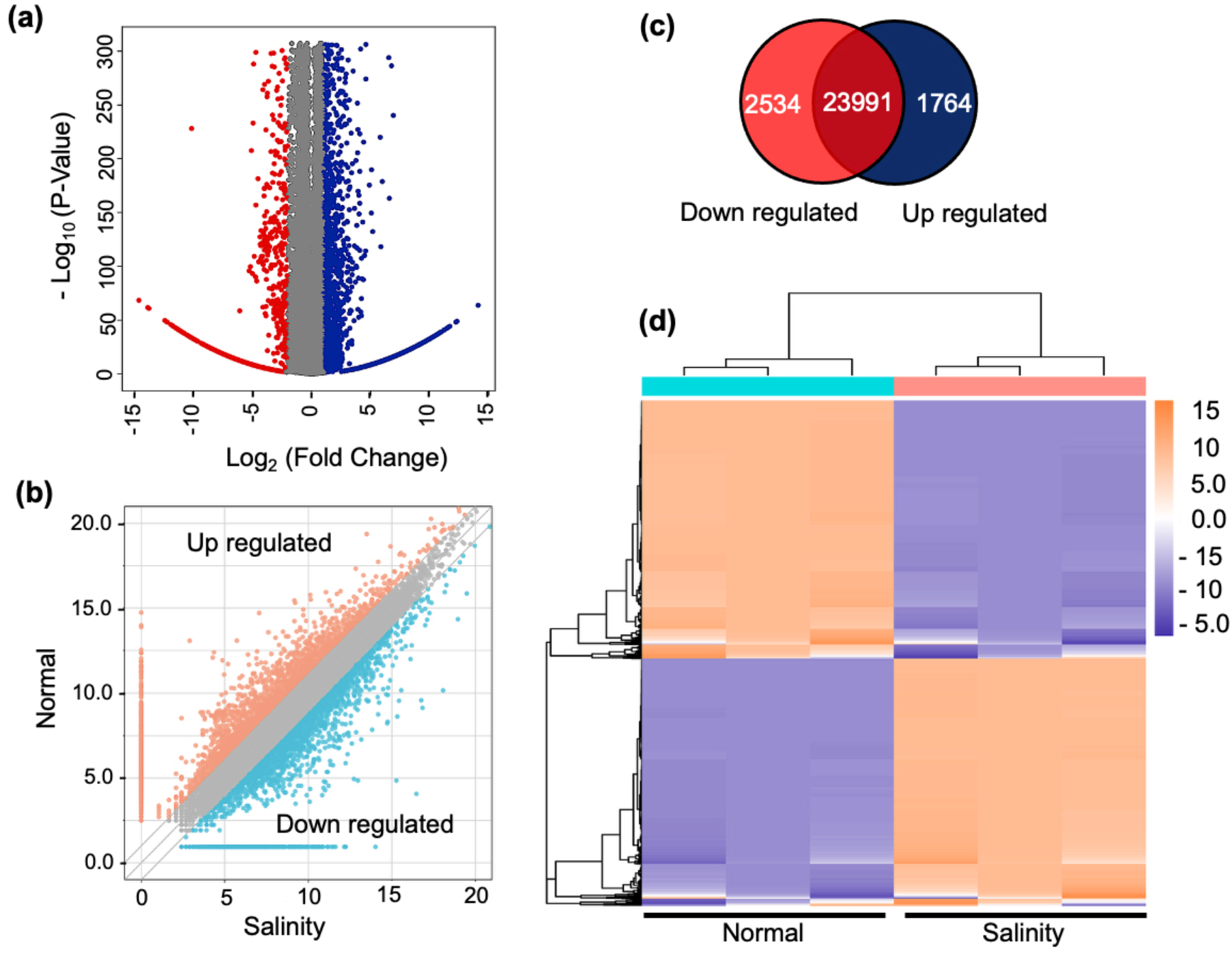
Transcriptomic analysis of differential gene expression in Giza 134 leaf blade under salinity stress. Differentially expressed genes (DEGs) were identified by comparing leaf blade tissues under normal and salinity stress conditions after 15 days of treatment. **(A)** Volcano plot showing the distribution of DEGs. The x-axis represents log_2_ fold change, and the y-axis represents –log₁₀(p-value). Vertical dotted lines indicate the 1.5-fold change threshold, and the horizontal dotted line marks the significance cutoff (*P* = 0.05). Blue dots represent significantly upregulated genes, and red dots indicate significantly downregulated genes. **(B)** MA plot displaying the relationship between mean expression (A) and log_2_ fold change (M) for each gene. Each dot represents an annotated gene, with DEGs highlighted based on differential expression between control and stress conditions. **(C)** Venn diagram showing the number of DEGs uniquely or commonly expressed between the normal and salinity-stressed groups. **(D)** Hierarchical clustering heatmap of DEGs under normal and salinity stress conditions. Rows represent genes, and columns represent biological replicates or conditions, with color gradients indicating relative expression levels.

These results indicate that Giza 134 exhibits extensive transcriptomic changes in response to salinity stress, involving the induction of stress-responsive genes and the repression of genes associated with growth and development. The identified differentially expressed genes provide a comprehensive dataset for subsequent functional and regulatory analyses.

### 3.6. Gene ontology enrichment analysis of DEGs

To elucidate the functional implications of salinity-induced transcriptional changes, Gene Ontology (GO) enrichment analysis was performed for upregulated and downregulated DEGs (FDR ≤ 0.05; Table S2 and S3). A total of 139 GO terms were significantly enriched among upregulated genes, whereas 124 terms were enriched among downregulated genes, spanning the categories of biological process (BP), molecular function (MF), and cellular component (CC).

Among the upregulated genes, enriched MF categories included hydrolase activity acting on glycosyl bonds (GO:0016798), O-glycosyl compound hydrolysis (GO:0004553), oxidoreductase activity (GO:0016702, GO:0016620), lipid binding (GO:0008289), and dioxygenase activity (GO:0051213) (Figure 5a). These functions are indicative of enhanced carbohydrate turnover, lipid-associated processes, and redox regulation, which are likely essential for osmoprotection, energy redistribution, and mitigation of oxidative stress under saline conditions. In the CC category, enrichment of terms such as extracellular region (GO:0005576), anchored component of membrane (GO:0031225), and plasma membrane (GO:0046658) underscores the importance of membrane-associated and secreted proteins in stress perception, signaling, and ion transport. At the BP level, significant enrichment of carbohydrate metabolism (GO:0005975), lipid metabolism (GO:0006629), and lipid transport (GO:0006869) further supports active metabolic reprogramming to maintain cellular homeostasis under salinity stress.

**Figure 5.**
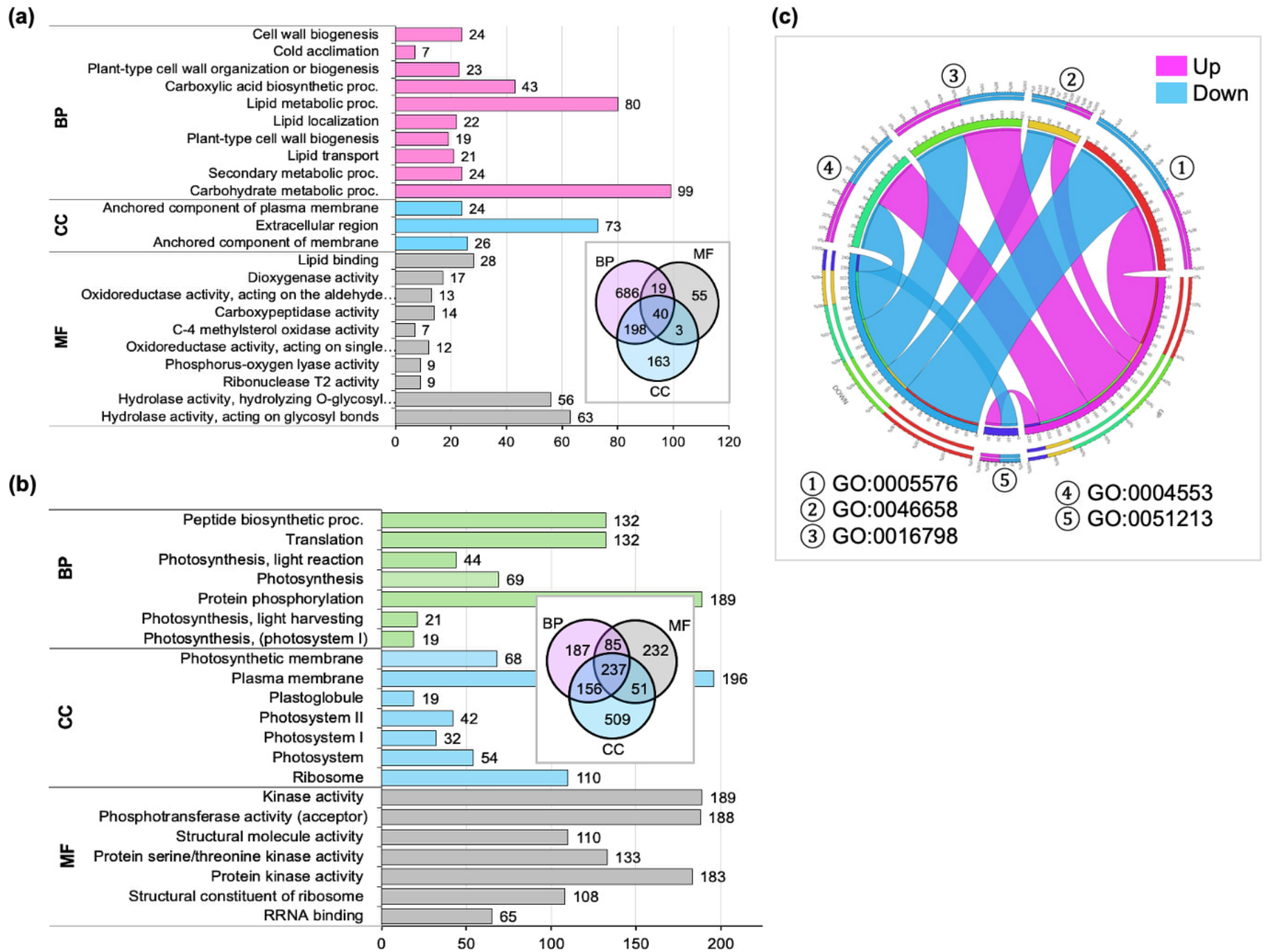
Transcriptomic response of barley to prolonged salinity stress. Gene Ontology (GO) enrichment analysis of significantly **(A)** up- and (B) down-regulated differentially expressed genes (DEGs) under prolonged salinity stress conditions (*P* ≤ 0.05). DEGs were defined based on a fold change threshold of Log_2_(FC) > 1 and a false discovery rate (FDR) < 0.05. GO terms were classified into three categories: Biological Process (BP), Cellular Component (CC), and Molecular Function (MF). The Venn diagram illustrates the overlap of enriched GO terms among the three categories. (C) Circos plot of common GO terms in up-regulated (blue sectors) and down-regulated (red sectors) genes under salinity stress. Each sector represents a GO term, and the connecting links indicate the overlap. The width of each link is proportional to the total number of genes associated with the GO term.

In contrast, downregulated genes were significantly enriched for MF categories related to structural molecule activity (GO:0005198), rRNA binding (GO:0019843), and kinase-associated functions, including protein kinase activity (GO:0004672), serine/threonine kinase activity (GO:0004674), and phosphotransferase activity (GO:0016773) (Figure 5b). These patterns suggest a reduction in protein synthesis and attenuation of signal transduction pathways, potentially reflecting an energy-conserving strategy under stress. Consistently, CC enrichment analysis revealed strong representation of *ribosome*-related terms (GO:0005840), *photosystems I* and *II* (GO:0009521, GO:0009522, GO:0009523), *photosynthetic membranes* (GO:0034357), *plastoglobules* (GO:0010287), and the *plasma membrane* (GO:0005886), indicating widespread downregulation of photosynthetic and translational machinery. At the BP level, repression of *photosynthesis* (GO:0015979), including *light reactions* (GO:0019684), light harvesting (GO:0009765), and *photosystem I activity* (GO:0009768), was evident, consistent with a suppression of energy production to limit photooxidative damage. Additionally, processes related to *translation* (GO:0006412), *peptide biosynthesis* (GO:0043043), and *protein phosphorylation* (GO:0006468) were significantly downregulated, highlighting a general shift away from growth-related functions.

Several GO terms were significantly enriched in both up- and downregulated gene sets, indicating that core functional categories are broadly affected but differentially regulated under salinity stress. Shared GO terms included extracellular region (GO:0005576), anchored component of the plasma membrane (GO:0046658), hydrolase activity acting on glycosyl bonds (GO:0016798), O-glycosyl compound hydrolysis (GO:0004553), and dioxygenase activity (GO:0051213) (Figure 5c; Table S2 and S3). This overlap highlights extensive remodeling of carbohydrate metabolism, cell wall–associated processes, and redox-related enzymatic activities, with regulatory direction likely determined by specific physiological demands during stress adaptation.

### 3.7. KEGG pathway enrichment analysis of DEGs

To further characterize the biological functions associated with salinity-responsive transcriptional changes, KEGG pathway enrichment analysis was performed. Nine significantly enriched pathways were identified among upregulated genes, while five pathways were significantly enriched among downregulated genes (Table 1).

**Table 1.** KEGG pathway classification of the DEGs, up-regulated and down-regulated in barley Giza 134 under salinity stress.

| DEGs | KEGG Pathway | Description | Intersection size | Term size | *p-value* | FDR |
| --- | --- | --- | --- | --- | --- | --- |
| Upregulated | ko00073 | Cutin, suberine and wax biosynthesis | 8 | 27 | 4.53E-07 | 0.0008 |
|  | ko00941 | Flavonoid biosynthesis | 9 | 36 | 1.08E-06 | 0.0008 |
|  | ko00591 | Linoleic acid metabolism | 6 | 16 | 1.36E-06 | 0.0008 |
|  | ko00500 | Starch and sucrose metabolism | 19 | 143 | 5.64E-06 | 0.0025 |
|  | ko00260 | Glycine, serine and threonine metabolism | 11 | 62 | 1.70E-05 | 0.0061 |
|  | ko00250 | Alanine, aspartate and glutamate metabolism | 9 | 49 | 8.28E-05 | 0.0213 |
|  | ko00130 | Ubiquinone and other terpenoid-quinone biosynthesis | 8 | 43 | 0.00021 | 0.0468 |
|  | ko00940 | Phenylpropanoid biosynthesis | 25 | 261 | 0.00027 | 0.0492 |
|  | ko00680 | Methane metabolism | 9 | 54 | 0.00027 | 0.0492 |
| Downregulated | ko00196 | Photosynthesis - antenna proteins | 21 | 31 | 2.63E-24 | 4.44E-21 |
|  | ko00194 | Photosynthesis proteins | 47 | 174 | 3.03E-21 | 2.56E-18 |
|  | ko03010 | Ribosome | 86 | 491 | 2.09E-19 | 1.18E-16 |
|  | ko03011 | Ribosome | 86 | 505 | 2.02E-18 | 8.54E-16 |
|  | ko00195 | Photosynthesis | 26 | 142 | 4.79E-07 | 1.62E-04 |
|  | ko00904 | Diterpenoid biosynthesis | 7 | 19 | 4.33E-05 | 1.22E-02 |

Consistent with the GO enrichment results, upregulated genes were predominantly associated with pathways involved in stress response and metabolic adjustment, including cutin, suberine, and wax biosynthesis (ko00073), flavonoid biosynthesis (ko00941), linoleic acid metabolism (ko00591), and starch and sucrose metabolism (ko00500). Additional enrichment was observed for amino acid metabolism pathways, such as glycine, serine, and threonine metabolism (ko00260) and alanine, aspartate, and glutamate metabolism (ko00250), as well as for pathways related to ubiquinone and terpenoid-quinone biosynthesis (ko00130) and phenylpropanoid biosynthesis (ko00940).

In contrast, downregulated genes were significantly enriched in pathways related to growth, energy production, and protein synthesis, including photosynthesis (ko00195), photosynthetic antenna proteins (ko00196), photosynthetic proteins (ko00194), and ribosome-associated pathways (ko03010 and ko03011). In addition, diterpenoid biosynthesis (ko00904) was significantly enriched among downregulated genes.

Collectively, KEGG pathway enrichment analysis indicates a clear functional divergence between upregulated and downregulated gene sets under salinity stress. Upregulated pathways are primarily associated with stress adaptation, metabolic adjustment, and protective functions, including reinforcement of surface barriers, antioxidant production, osmoprotective metabolism, and secondary metabolite biosynthesis. In contrast, downregulated pathways are predominantly linked to photosynthesis, energy production, and protein synthesis, reflecting a coordinated reduction in growth- and energy-demanding processes under saline conditions.

### 3.8. Candidate genes for salinity tolerance

Salinity stress was associated with marked changes in the expression of genes involved in stress perception, signal transduction, osmoprotection, metabolic processes, and redox regulation (Table 2). Among the most strongly upregulated genes in Giza 134, several are known to be responsive to salinity and other abiotic stresses. These included *HORVU6Hr1G060810*, encoding a serine/threonine protein kinase, and the dehydrin genes DHN4 (*HORVU6Hr1G083980*) and DHN3 (*HORVU6Hr1G084070*). Dehydrins are classical markers of abiotic stress and play critical roles in osmotic adjustment, membrane stabilization, and protection of proteins during dehydration and salinity stress (Abedini et al. 2017; Close 2006). Their strong induction indicates that osmotic stress represents a dominant component of the salinity response in the Giza 134 cultivar. In addition, ascorbate peroxidase 1 (APX1) (*HORVU6Hr1G088460*) showed strong induction under salinity stress.

**Table 2.**
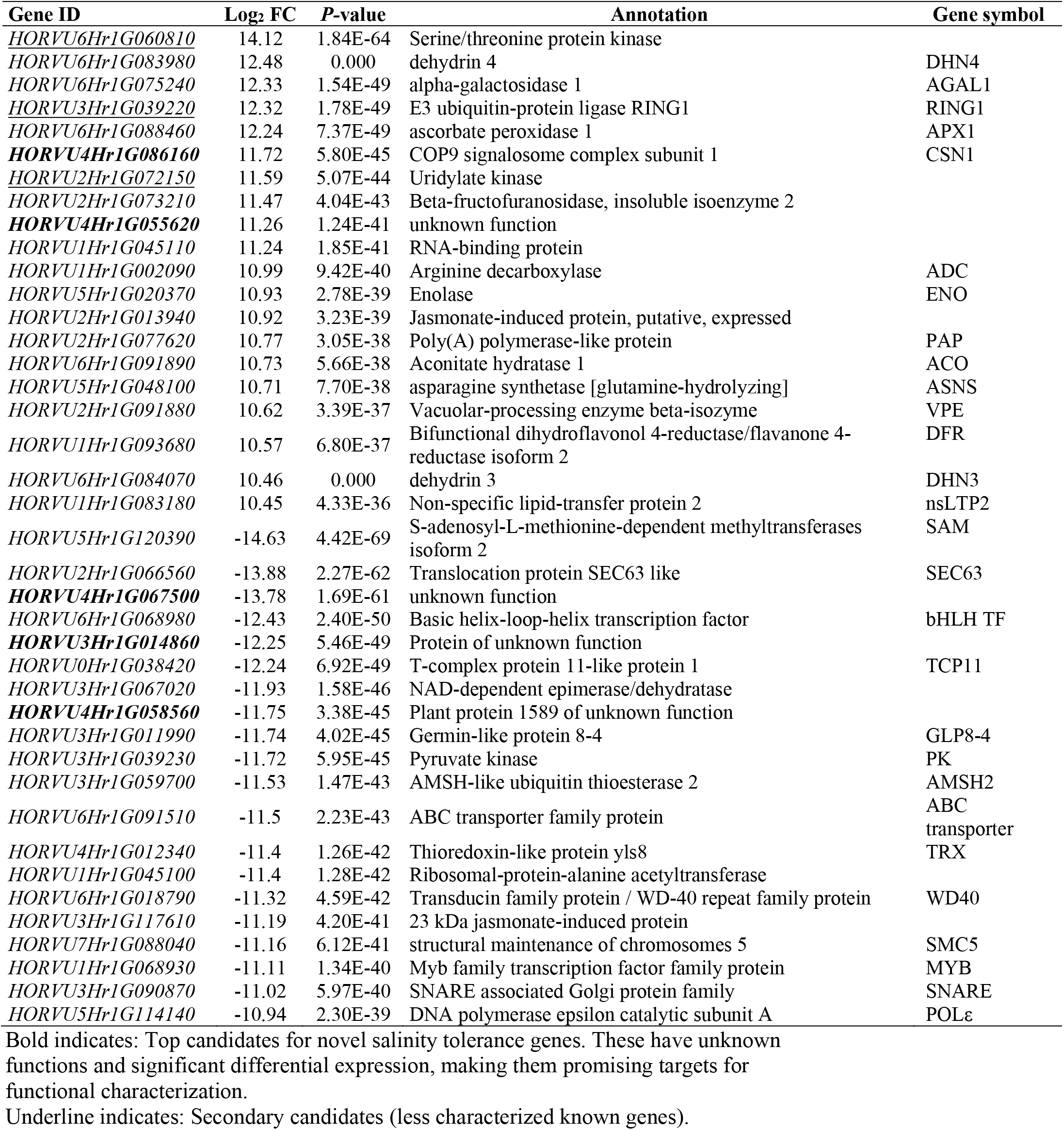
Functional annotations of the top 20 DE genes, up- and down-regulated in response to salinity stress.

Other upregulated genes included E3 ubiquitin-protein ligase RING1 (*HORVU3Hr1G039220*), arginine decarboxylase (ADC) (*HORVU1Hr1G002090*), and enzymes associated with central carbon metabolism, such as enolase (ENO) (*HORVU5Hr1G020370*) and aconitate hydratase (ACO) (*HORVU6Hr1G091890*). A poly(A) polymerase–like protein (PAP) (*HORVU2Hr1G077620*) was also among the highly induced genes.

In contrast, several TFs and core cellular regulators were strongly downregulated under salinity stress. These included a basic helix–loop–helix (bHLH) transcription factor (*HORVU6Hr1G068980*) and a MYB transcription factor (*HORVU1Hr1G068930*). Genes involved in DNA replication and genome maintenance, such as DNA polymerase epsilon catalytic subunit (POLε) (*HORVU5Hr1G114140*) and structural maintenance of chromosomes protein 5 (SMC5) (*HORVU7Hr1G088040*), were also downregulated.

In addition to well-annotated genes, several genes with limited or unknown functional annotation exhibited strong differential expression. COP9 signalosome subunit 1 (CSN1) (*HORVU4Hr1G086160*) was among the highly upregulated genes, whereas *HORVU4Hr1G055620* (unknown function) ranked among the most induced candidates. Conversely, *HORVU4Hr1G067500*, an uncharacterized gene, showed strong downregulation under salinity stress.

Tissue-specific expression profiling of selected candidate genes revealed distinct expression patterns across barley tissues (Figure 6). Several upregulated genes, including dehydrins and ROS-scavenging enzymes, showed higher expression in roots, whereas many downregulated TFs and cell cycle–related genes were predominantly expressed in reproductive and meristematic tissues and developing grains (e.g., CAR5, CAR15).

**Figure 6.**
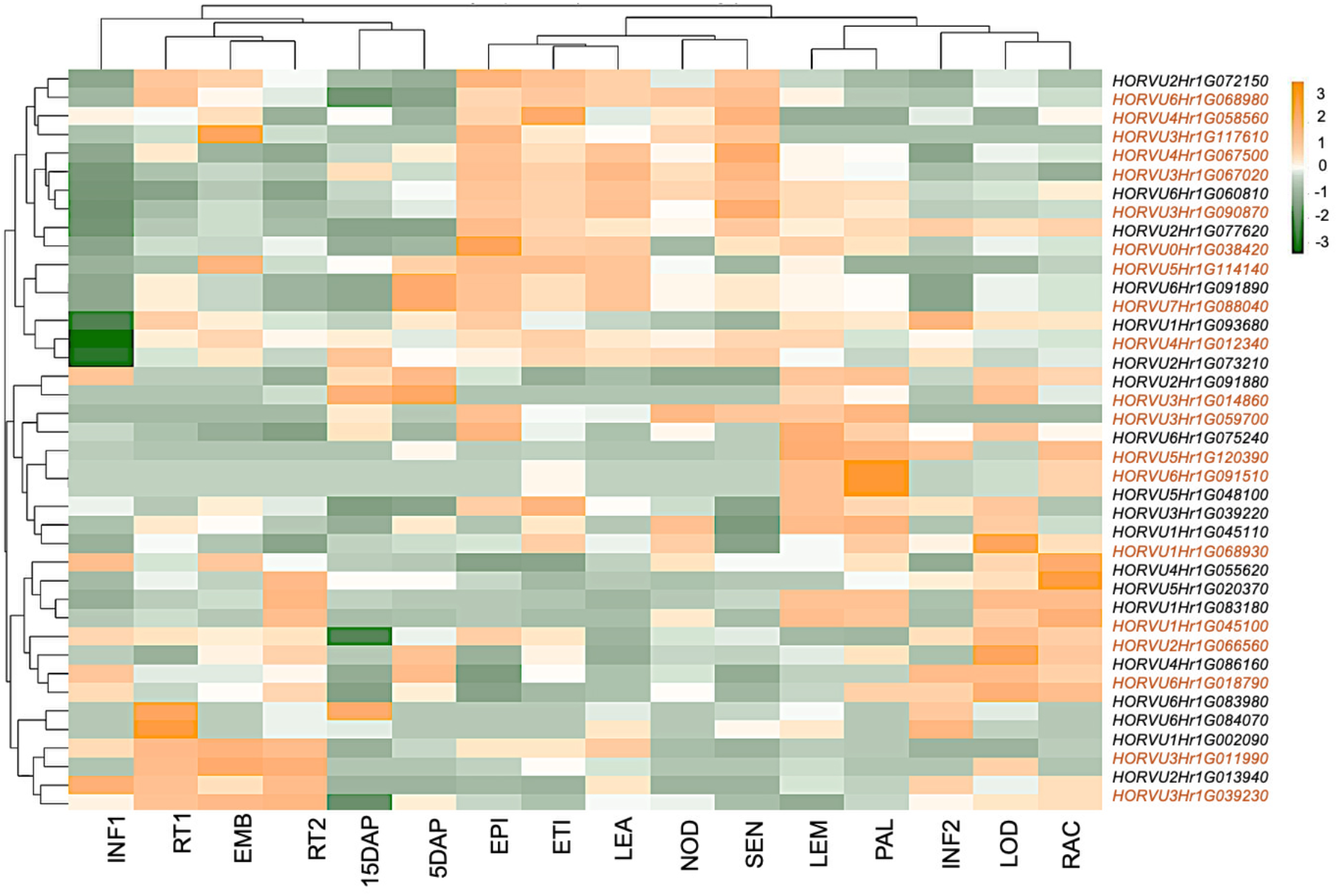
Expression profiles of the 40 candidate genes across diverse developmental stages and tissues. Heatmap depicting the expression levels of 40 candidate genes (y-axis) across 16 barley tissues and developmental stages (x-axis). Expression values are visualized as z-scores of normalized transcripts per million (TPM) to highlight tissue-specific expression patterns relative to the mean for each gene (see colour scale). Gene identifiers for the 40 candidates are listed. Expression data were retrieved and analyzed from the BarleyNBI (www.inetbio.org/barleynet/) platform. EMB: 4-day embryos, ROO1: Roots from seedlings (10 cm shoot stage), LEA: Shoots from seedlings (10 cm shoot stage), INF1: Young developing inflorescences (5 mm), INF2: Developing inflorescences (1-1.5 cm), NOD: Developing tillers, 3rd internode (42 DAP), CAR5: Developing grain (5 DAP), CAR15: Developing grain (15 DAP), ETI: Etiolated seedling, dark cond. (10 DAP), LEM: Inflorescences, lemma (42 DAP), LOD: Inflorescences, lodicule (42 DAP), PAL: Dissected inflorescences, palea (42 DAP), EPI: Epidermal strips (28 DAP), RAC: Inflorescences, rachis (35 DAP), ROO2: Roots (28 DAP), SEN: Senescing leaves (56 DAP)

Taken together, these results highlight a diverse set of salinity-responsive candidate genes in Giza 134, including signaling kinases, TFs, metabolic enzymes, stress-associated proteins such as dehydrins and peroxidases, and components of protein turnover pathways. These genes represent a comprehensive resource for subsequent functional and regulatory analyses.

### 3.9. *Cis-*regulatory motif analysis under salinity stress

To characterize transcriptional regulatory features associated with salinity stress, *cis*-regulatory motif enrichment analysis was performed on the promoter regions of DEGs. Distinct motif enrichment patterns were observed between upregulated and downregulated gene sets, indicating contrasting regulatory signatures under salinity stress.

Among upregulated genes, significant enrichment was detected for motifs associated with MYB-related TFs (e.g., ACCCTA, CTTCTC), IDD (CAAAAC), G2-like (AAGGAA), and MADS-box TFs (Figure 7a; Table S4). These motifs were consistently overrepresented in promoter regions of genes induced under salinity stress.

**Figure 7.**
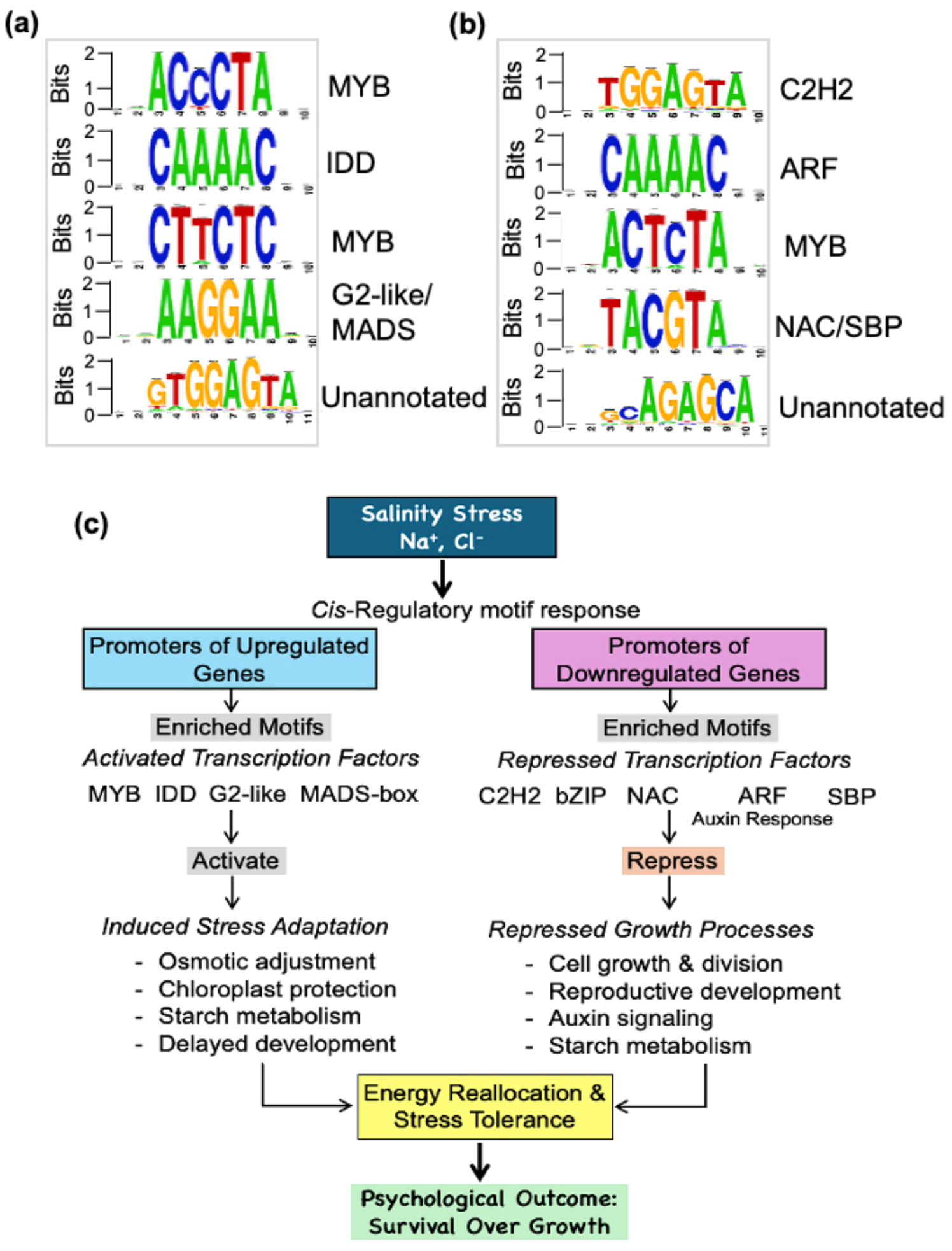
Analysis of transcription factor binding sites in the promoter of differentially expressed genes. **(A)** The sequence logos for TFs whose binding sites were significantly enriched in the promoter of up-regulated genes. **(B)** The sequence logos for TFs whose binding sites were significantly enriched in the promoter of down-regulated genes. **(C)** A model of transcriptional reprogramming in spring barley under salinity stress in this study. The influx of salt ions triggers a shift in gene expression through the enrichment of specific *cis*-regulatory motifs in the promoters of differentially expressed genes. This leads to the activation of stress-adaptive pathways (blue) via TFs like MYB, IDD, G2-like, and MADS-box, and the repression of growth-related pathways (red) via TFs like Zinc finger, ARF, bZIP, NAC, and SBP. This dual mechanism allows the plant to prioritize energy towards tolerance and survival.

In contrast, promoters of downregulated genes were significantly enriched for motifs corresponding to zinc finger (C2H2) proteins (TGGAGTA), auxin response factors (ARFs; CAAAAC), MYB-related telomere-binding factors (ACTCTA), bZIP TFs (TACGTA), and members of the NAC and SBP transcription factor families (Figure 7b; Table S4).

In addition to known motifs, two previously unannotated sequence motifs were identified: GTGGAGTA, which was enriched among upregulated genes, and AGAGCA, which was enriched among downregulated genes. These motifs showed significant positional enrichment within promoter regions and may represent additional cis-regulatory elements associated with salinity-responsive gene expression.

Overall, *cis*-regulatory motif analysis reveals a dual transcriptional regulatory strategy in barley under salinity stress: (i) activation of stress-adaptive gene expression programs mediated by MYB, IDD, G2-like, and MADS-box TFs, and (ii) repression of growth-and reproduction-associated pathways through zinc finger, ARF, NAC, bZIP, and SBP TFs (Figure 7c). This coordinated transcriptional reprogramming reflects a strategic balance between mitigating salt-induced damage and maintaining essential physiological functions necessary for survival under saline conditions.

### 3.10. Validation of the DEGs expression using RT-qPCR

To validate the reliability of the RNA-seq data, the expression patterns of three representative differentially expressed genes were examined using RT-qPCR (Figure 8). Two genes identified as upregulated under salinity stress in the RNA-seq analysis, Serine/threonine protein kinase (*HORVU6Hr1G060810*) and Dehydrin 4 (*HORVU6Hr1G083980*), showed strong induction under salinity conditions, with transcript levels increasing approximately 7.65-fold and 9.07-fold, respectively, compared with the control. In contrast, the expression of Ribosomal protein alanine acetyltransferase (*HORVU1Hr1G045100*), which was downregulated in the RNA-seq dataset, was significantly reduced under salinity stress to approximately 45% of the control level.

**Figure 8.**
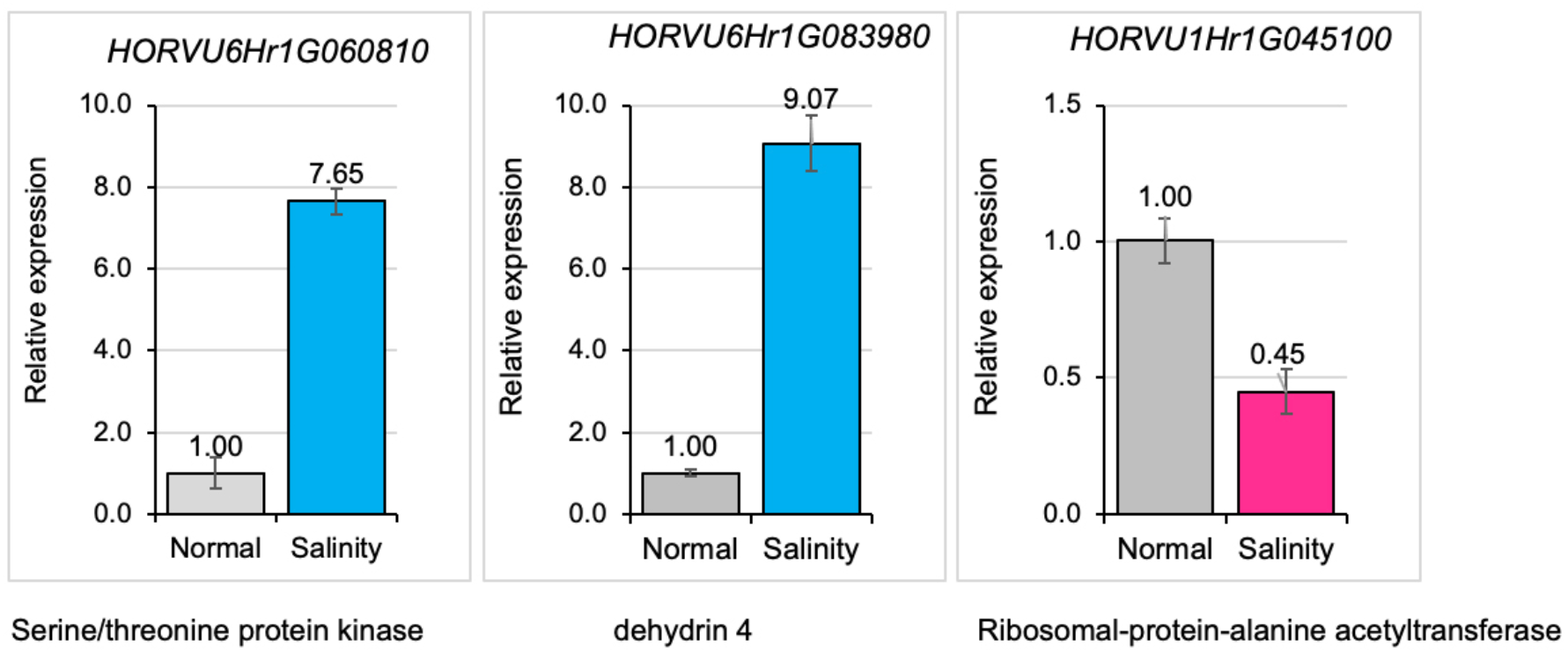
RT-qPCR validation of RNA-seq results. Relative expression levels of three representative differentially expressed genes under Normal and salinity stress conditions. Two genes, Serine/threonine protein kinase (*HORVU6Hr1G060810*) and Dehydrin 4 (*HORVU6Hr1G083980*), were upregulated under salinity stress, whereas Ribosomal protein alanine acetyltransferase (*HORVU1Hr1G045100*) was downregulated. Expression levels were normalized to an internal reference gene and are presented relative to the control. Fold changes are indicated above the bars. Error bars indicate mean ± SD of biological replicates.

Overall, the RT-qPCR results were highly consistent with the RNA-seq data in both the direction and magnitude of gene expression changes, confirming the robustness and accuracy of the transcriptomic analysis.

## 4. Discussion

In this study, the field, lysimeter, and growth chamber experiments were designed to address complementary aspects of salinity stress in barley. The field and lysimeter experiments evaluated physiological, biochemical, and agronomic responses, whereas the growth chamber experiment characterized transcriptomic responses under controlled conditions. The findings from these experiments were integrated conceptually to provide a comprehensive understanding of salinity responses and were not intended for direct statistical comparison.

### 4.1. Salinity stress severely constrains growth and yield through physiological and biochemical disruption

Salinity stress significantly impaired growth performance and agronomic productivity in the barley cultivar Giza 134, as reflected by significant reductions in plant height, biological yield, grain yield, and 1000-grain weight. These reductions were evident under both field salinity (SS1) and lysimeter-based salinity stress (SS2), with the latter imposing a stronger inhibitory effect (Figure 1). The more pronounced yield penalties observed under SS2 likely reflect the higher uniformity and persistence of salt exposure in the lysimeter system. Similar growth suppression under saline conditions has been widely reported in barley and other cereals, where osmotic stress and ion toxicity restrict cell expansion, disrupt metabolic processes, and limit assimilate partitioning to reproductive organs (Elakhdar et al. 2016; Gupta and Huang 2014; Munns and Tester 2008). The observed decline in LA and SLW under salinity stress further indicates structural and developmental constraints on leaf growth (Figure 2A). Reduced leaf expansion is a common adaptive response to osmotic stress, limiting transpirational water loss but simultaneously reducing photosynthetic surface area and carbon assimilation capacity (Chaves et al. 2009; Skirycz et al. 2011; Skirycz and Inze 2010). Consistent with these morphological changes, RWC decreased significantly under salinity stress, reflecting impaired plant water status and osmotic imbalance. RWC often indicate insufficient osmotic adjustment and reduced water uptake under saline conditions (Elakhdar et al. 2016; Gupta and Huang 2014; Negrao et al. 2017).

Photosynthetic pigment include, *chl a* and *chl b* declined in Giza 134 under saline conditions, resulting in a reduction in total chlorophyll content (Figure 2B). Notably, *chl b* showed a stronger decline than *chl a*, leading to a decrease in the chlorophyll a/b ratio. This shift suggests disruption of the light-harvesting complex and possible degradation of photosystem-associated pigments, which can impair light absorption efficiency and overall photosynthetic performance (Parida and Das 2005; Stepien and Johnson 2009). Chlorophyll degradation under salinity has been associated with oxidative damage to chloroplast membranes, inhibition of chlorophyll biosynthesis, and enhanced pigment breakdown triggered by ROS accumulation (Hammami et al. 2024; Munns and Tester 2008). In contrast, proline accumulated substantially in response to salinity stress (Figure 2B). Proline is widely recognized as a multifunctional osmoprotectant that contributes to osmotic adjustment, ROS detoxification, and stabilization of cellular structures under stress conditions (Arias-Baldrich et al. 2015; Ashraf and Foolad 2007; Ben Rejeb et al. 2014; Szabados and Savoure 2010). Thus, strong induction of proline observed indicates activation of protective metabolic pathways in Giza 134 aimed at maintaining cellular homeostasis under saline environments. However, the persistence of significant growth and yield reductions suggests that proline accumulation alone is insufficient to fully mitigate the damaging effects of prolonged salt exposure (Lehmann et al. 2010; Szabados and Savoure 2010).

Salinity stress also disrupted ionic homeostasis, as evidenced by the significant accumulation of Na^+^ and the concomitant reduction in K^+^ concentration, resulting in a significantly elevated Na^+^/K^+^ ratio (Figure 3A). Maintenance of a low Na^+^/K^+^ ratio is essential for enzymatic activity, membrane stability, and metabolic regulation (Chen et al. 2007; Shabala and Cuin 2008; Zhu et al. 2015). The ionic imbalance induced by salinity was accompanied by increased lipid peroxidation, as indicated by elevated MDA levels. In response to this oxidative challenge, antioxidant defense mechanisms were activated, as demonstrated by the significant increases in CAT and POD activities (Figure 3B). These enzymes play critical roles in detoxifying hydrogen peroxide and other ROS, thereby protecting cellular components from oxidative injury. The upregulation of antioxidant enzymes observed here suggests that Giza 134 initiates an enzymatic defense response to mitigate oxidative stress; however, the persistence of elevated MDA levels indicates that ROS generation under high salinity may exceed the detoxification capacity of the antioxidant system (Apel and Hirt 2004; Gill and Tuteja 2010b; Mittler 2002).

These results demonstrate that salinity stress imposes a complex cascade of physiological and biochemical disturbances in Giza 134, including impaired water status, reduced photosynthetic capacity, ionic imbalance, and oxidative stress, ultimately lead to substantial reductions in plant growth and grain productivity, particularly under severe and sustained salinity conditions.

### 4.2. Extensive transcriptomic reprogramming under salinity stress reflects a shift from growth to survival

RNA-seq analysis revealed extensive transcriptional reprogramming in response to salinity stress, with more than 4,000 genes differentially expressed. The predominance of downregulated genes suggests a broad suppression of growth-related processes, accompanied by selective activation of stress-responsive pathways, reflecting a conserved response in which cellular resources are redirected from growth and development toward stress mitigation (Deinlein et al. 2014; Gharaghanipor et al. 2022; Ziemann et al. 2013).

### 4.3. Metabolic and antioxidative pathways support salinity adaptation

Functional enrichment analyses revealed that genes upregulated under salinity stress were primarily associated with carbohydrate metabolism, lipid metabolism, oxidoreductase activity, and secondary metabolite biosynthesis, indicating substantial metabolic reprogramming in response to salt-induced physiological disturbances (Figure 5). These transcriptional changes are consistent with the physiological responses observed in this study (Figure 1-3), including reduced water status, ionic imbalance, and increased oxidative stress, all of which require coordinated metabolic adjustments to maintain cellular homeostasis (Du and Benning 2016; Gill and Tuteja 2010b; Gupta and Huang 2014).

Enhanced expression of genes involved in starch and sucrose metabolism (Figure 5A), suggests activation of carbohydrate mobilization pathways that may contribute to osmotic adjustment under saline conditions, as soluble sugars maintain cellular osmotic balance and protect macromolecules during dehydration stress (Keunen et al. 2013; Rasmusson et al. 2009). This response likely complements the substantial accumulation of proline observed in Giza 134, together supporting osmotic stabilization when salinity-induced osmotic stress reduces water uptake and relative water content.

Lipid and fatty acid metabolism pathways were also strongly enriched among upregulated genes (Figure 5A). Membrane lipid remodeling is a well-established adaptive mechanism may help maintain membrane fluidity and structural integrity under stress while generating lipid-derived signaling molecules that regulate stress responses (Du and Benning 2016; Hou et al. 2016; Upchurch 2008). Such membrane adjustments are particularly important under saline conditions, where ionic toxicity and oxidative stress can destabilize cellular membranes, as reflected by the increased MDA observed in Giza 134. Consistent with this, the enrichment of cuticular wax biosynthesis pathways suggests additional structural adaptations that may reduce transpirational water loss and enhance tolerance to salt-induced dehydration (Zhu and Xiong 2013). In parallel, pathways associated with flavonoid and phenylpropanoid biosynthesis were significantly enriched among upregulated genes (Table 1). Flavonoids are important secondary metabolites that function as non-enzymatic antioxidants capable of scavenging ROS and protecting cellular components from oxidative damage (Nakabayashi et al. 2014). The transcriptional activation of these pathways is therefore consistent with the elevated oxidative stress indicated by increased MDA levels and the induction of antioxidant enzymes such as CAT and POD (Figure 3B). Enhanced flavonoid production has been widely associated with improved tolerance to salinity and other abiotic stresses (Hou et al. 2016; Jan et al. 2021).

Additionally, the induction of amino acid metabolism pathways, including those involving glycine, serine, threonine, and glutamate (Table 1), likely supports the synthesis of compatible solutes and stress-related metabolites. These pathways may also facilitate nitrogen remobilization and metabolic flexibility during stress adaptation (Batista-Silva et al. 2019; Hildebrandt et al. 2015; Krasensky and Jonak 2012). In particular, amino acid metabolism is closely linked to the biosynthesis of osmoprotectants such as proline, which accumulated strongly in Giza 134 under saline conditions.

### 4.4. Candidate genes suggest potential regulators of stress signaling and cellular protection

Analysis of highly responsive genes identified several components associated with salinity tolerance (Table 2). Strong induction of dehydrins (*DHN3* and *DHN4*) is consistent with a role in osmotic protection and membrane stabilization under salt stress (Abedini et al. 2017; Close 2006). Upregulation of ascorbate peroxidase (*APX1*) is consistent with the involvement of ROS detoxification via the ascorbate-glutathione cycle (Mittler 2002; Shigeoka et al. 2002). Salinity stress was associated with increased expression of genes involved in central carbon and nitrogen metabolism, including enolase, aconitate hydratase, asparagine synthetase, and arginine decarboxylase, suggesting metabolic flexibility as a key component of stress adaptation (Alcazar et al. 2010; Foyer and Noctor 2005; Gill and Tuteja 2010a). In parallel, induction of signaling and protein turnover components, such as *serine/threonine kinases*, *RING-type E3 ubiquitin ligases*, and *COP9 signalosome subunit 1*, indicates tight regulatory control of stress signaling and protein stability (Hunter 2007; Lyzenga and Stone 2012; Vierstra 2009; Wei et al. 2008).

Conversely, strong repression of genes associated with DNA replication, chromatin maintenance, protein trafficking, and transcriptional regulation reflects stress-induced growth restraint (Table 2). Downregulation of *methyltransferases*, *DNA polymerase ε*, *SMC5*, and vesicle trafficking components (*SEC63* and *SNARE*) suggests coordinated suppression of cell cycle progression and cellular maintenance processes under salinity stress (Howell 2013; Serrano-Mislata et al. 2025; Skirycz and Inze 2010). Additionally, the repression of TFs, such as *bHLH* and *MYB* family proteins, supports large-scale transcriptional reprogramming, with developmental and growth-related gene networks being downregulated in favor of stress defense pathways (Dubos et al. 2010; Pires and Dolan 2010).

### 4.5. *Cis-*regulatory motifs reveal coordinated transcriptional control

*Cis*-regulatory motif analysis revealed distinct promoter architectures between upregulated and downregulated genes, providing insight into transcriptional control under salinity stress (Figure 7). Enrichment of MYB-, IDD-, G2-like-, and MADS-box–related motifs among upregulated genes supports the involvement of transcriptional regulators associated with metabolic regulation, stress signaling, and developmental plasticity (Ambawat et al. 2013; Zhang et al. 2025). This pattern is consistent with findings in rice and *Arabidopsis*, where MYB and MADS-box TFs play central roles in regulating carbohydrate metabolism, chloroplast development, flowering time, and abiotic stress responses, including salinity and drought tolerance (Dubos et al. 2010; Roy 2016; Shinozaki and Yamaguchi-Shinozaki 2007; Smaczniak et al. 2012). In contrast, enrichment of zinc finger, ARF, NAC, bZIP, and SBP motifs among downregulated genes reflects repression of hormone-mediated signaling and growth-related transcriptional programs, consistent with stress-induced growth restraint observed in cereals such as wheat and maize (Forestan et al. 2020; Jing et al. 2023; Nakashima et al. 2009). The detection of previously unannotated motifs suggests the presence of additional regulatory elements that may contribute to species-specific or condition-dependent salinity responses in barley.

Overall, the cis-regulatory motif landscape supports a model suggesting that salinity stress is associated with induces a highly coordinated transcriptional response in barley through the selective activation and repression of distinct TF families. This multilayered regulatory framework likely underlies the dynamic balance between stress defense, metabolic adjustment, and growth suppression, enabling effective acclimation to saline environments.

### 4.6. Integrated model of salinity stress responses in Giza 134

Salinity stress is associated with a complex cascade of physiological, biochemical, and transcriptional responses in the barley cultivar Giza 134, as summarized in the integrated model proposed in Figure 9. Exposure to saline conditions initially imposes osmotic stress and ionic toxicity, primarily resulting from excessive Na^+^ accumulation and disruption of the cellular Na^+^/K^+^ balance (Figure 3A). These disturbances impair water uptake and cellular hydration, leading to reduced RWC and inhibition of photosynthetic processes, as reflected by the decline in chlorophyll pigments (Figure 2). Impaired photosynthetic activity promotes the generation of reactive ROS, which act both as damaging agents and signaling molecules. Elevated ROS levels cause oxidative damage to membrane lipids, indicated by increased MDA accumulation. In response to this oxidative challenge, Giza 134 activates multiple protective mechanisms to maintain cellular homeostasis. The enzymatic antioxidant system, including CAT and POD, is enhanced to detoxify ROS and protect cellular components from oxidative injury. In parallel, the accumulation of compatible solutes such as proline contributes to osmotic adjustment and stabilization of proteins and membranes under saline conditions. The induction of proline biosynthesis pathways, particularly through key enzymes such as *P5CS* and *PSS*, further reflects activation of osmoprotective mechanisms.

**Figure 9.**
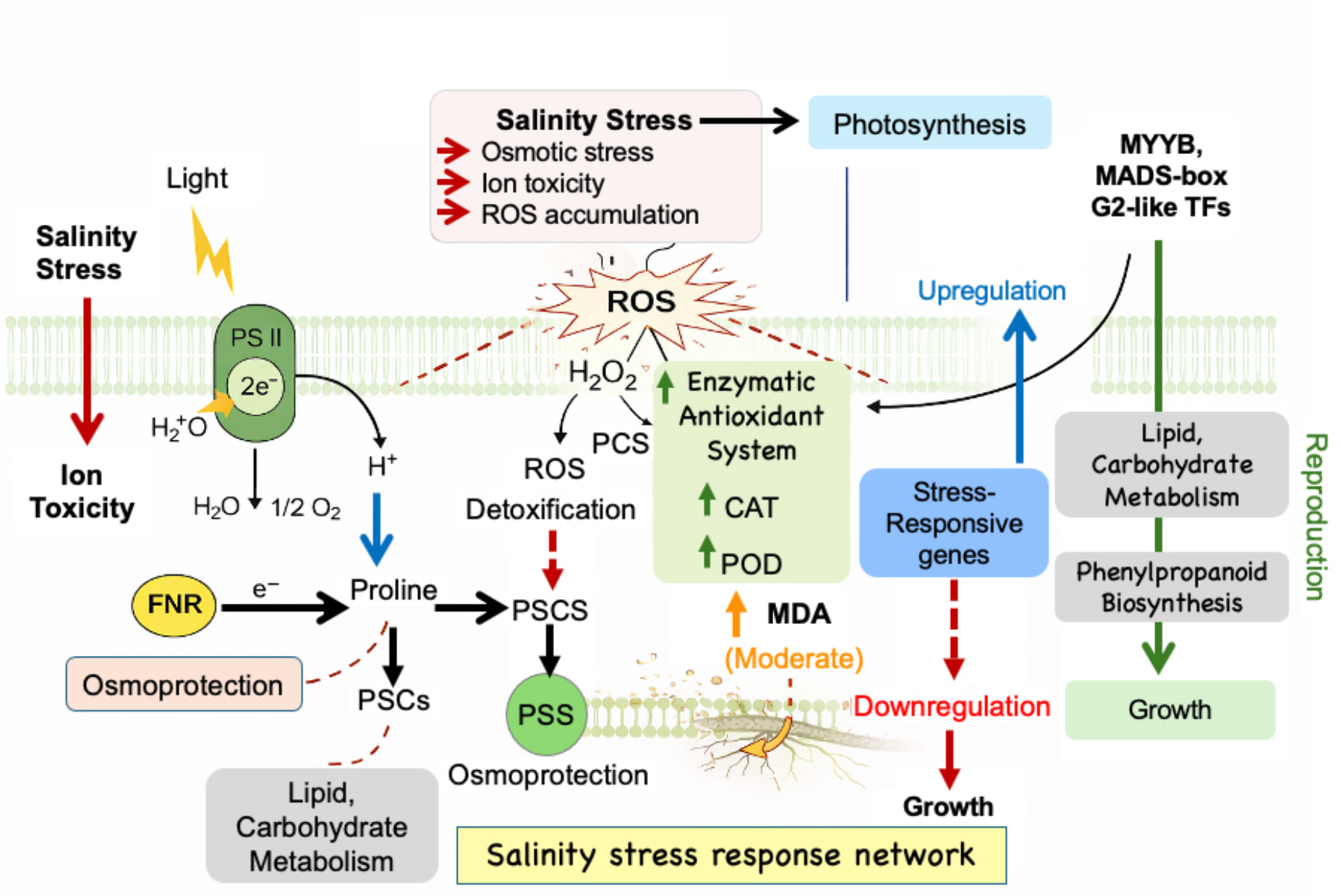
Proposed regulatory network underlying salinity stress response in spring barley (cv. Giza 134). Salinity stress induces osmotic imbalance and ion toxicity, leading to disruption of photosynthetic electron transport and enhanced production of ROS, particularly H_2_O_2_. Transcriptomic analysis revealed upregulation of stress-responsive TFs (MYB, MADS-box, and G2-like) and activation of pathways associated with lipid remodeling, carbohydrate metabolism, and phenylpropanoid biosynthesis. Biochemical assays demonstrated significant increases in antioxidant enzyme activities, including catalase (CAT), and peroxidase (POD), indicating activation of the enzymatic ROS-scavenging system. Although malondialdehyde (MDA) levels increased under salt stress, the magnitude of accumulation was moderate, suggesting controlled lipid peroxidation and effective mitigation of oxidative damage. Proline biosynthesis (via P5CS and PSS) was enhanced, contributing to osmoprotection and cellular stabilization. Concurrently, ribosome biogenesis and growth-related processes were downregulated, reflecting a shift from biomass accumulation to stress defense. Collectively, the model highlights ROS-mediated signaling as a central hub coordinating transcriptional and metabolic reprogramming in Giza 134 under salinity stress.

Transcriptomic analysis also reveals activation of metabolic pathways involved in carbohydrate and lipid metabolism, phenylpropanoid biosynthesis, and amino acid metabolism (Figure 5 and Tbale 1). These pathways may contribute to osmotic balance, membrane stability, and antioxidant defense. Secondary metabolites, including flavonoids derived from phenylpropanoid metabolism, likely contribute to non-enzymatic ROS scavenging and enhance cellular protection. The regulation of these metabolic adjustments is coordinated by stress-responsive TFs, including members of the MYB, MADS-box, and G2-like families, are predicted to regulate or may regulate the expression stress tolerance pathways. At the same time, many growth-related processes are transcriptionally downregulated, reflecting a shift in resource allocation from growth and biomass production toward stress defense and survival. This integrated response network may contribute to the observed salinity response in Giza 134 to partially mitigate salinity-induced damage; however, prolonged exposure to high salinity ultimately leads to significant reductions in plant growth and yield.

## Acknowledgments

This work was supported by a Grant-in-Aid for Scientific Research from the Japan Society for the Promotion of Science, grant number 19F19394. Ammar Elakhdar is thankful to the Barley Research Department, Field Crops Research Institute of the Agricultural Research Center in Egypt for providing the seeds of the Giza 134 cultivar.

## Author contributions

A.E. developed the article concept, conceptualization, investigation, methodology performed bioinformatics, interpretation of data, wrote the original draft. E.A. morphology data collection, review & editing. D.E. physiological data collection, analysis, review & editing. T.K. review & editing, visualization, supervision. All authors have read and approved the final mauscript.

## Data availability statement

The data that supports the findings of this study are available in the supplementary material of this article.

